# Multi-dimensional and spatiotemporal correlative imaging at the plasma membrane of live cells to determine the continuum nano-to-micro scale lipid adaptation and collective motion

**DOI:** 10.1101/2021.03.02.433589

**Authors:** Miguel Bernabé-Rubio, Minerva Bosch-Fortea, Miguel A. Alonso, Jorge Bernardino de la Serna

**Affiliations:** Department of Cell Biology and Immunology, Centro de Biología Molecular Severo Ochoa, Consejo Superior de Investigaciones Científicas and Universidad Autónoma de Madrid, Madrid, 28049 Spain; King’s College London Centre for Stem Cells and Regenerative Medicine, 28th Floor, Tower Wing, Guy’s Campus, Great Maze Pond, London, SE1 9RT UK; Institute of Bioengineering and School of Engineering and Materials Science, Queen Mary, University of London, Mile End Road, London, E1 4NS UK; Central Laser Facility, Rutherford Appleton Laboratory, MRC-Research Complex at Harwell, Science and Technology Facilities Council, Harwell, OX11 0QX UK; National Heart and Lung Institute, Imperial College London, Sir Alexander Fleming Building, London, SW7 2AZ UK; NIHR Imperial Biomedical Research Centre, London, SW7 2AZ UK

**Author notes:** Equal contribution (Miguel Bernabé-Rubio, Minerva Bosch-Fortea).

## Abstract

The primary cilium is a specialized plasma membrane protrusion with important receptors for signalling pathways. In polarized epithelial cells, the primary cilium assembles after the midbody remnant (MBR) encounters the centrosome at the apical surface. The membrane surrounding the MBR, namely remnant associated membrane patch (RAMP) once situated next to the centrosome, releases some of its lipid components to form a centrosome-associated membrane patch (CAMP) from which the ciliary membrane stems. The RAMP undergoes a spatiotemporal membrane refinement during the formation of the CAMP, which becomes highly enriched in condensed membranes with low lateral mobility. To better understand this process, we have developed a correlative imaging approach that yields quantitative information about the lipid lateral packing, its mobility and collective assembly at the plasma membrane at different spatial scales over time. Our work paves the way towards a quantitative understanding of lipid collective assembly at the plasma membrane spatiotemporally as a functional determinant in cell biology and its direct correlation with the membrane physicochemical state. These findings allowed us to gain a deeper insight into the mechanisms behind the biogenesis of the ciliary membrane of polarized epithelial cells.

## Introduction

Understanding how complex cellular structures function at the molecular level employing imaging approaches requires further technological and methodological advancements. Often, these challenges require determining structure-function relationships, which more recently are being tackled by means of combinatorial microscopy^1^ (i.e., employing different instruments). Correlative microscopy applied to cells often refers to the combination of optical and electron microscopy techniques to resolve molecules labelled fluorescently and with an electro-dense marker at different spatial and temporal resolution. These multimodal approaches have gained popularity since they have the potential to disentangle the molecular or cellular ultrastructure and correlate it with the biological function. However, these techniques are limited to a particular timepoint in dynamic biological process. Other combinatorial imaging methods can reveal molecular events spatiotemporally at different nano- and micro- scales using, for instance, cross-correlating approaches. In optical fluorescence microscopy, these other correlative imaging modalities can also involve super-resolution microscopy, thus advantageously resolving functional events in a simultaneous manner when they happen as they happen. Although, these methods cannot resolve ultrastructure architecture to the same detail, they can provide clues towards revealing molecular structure-function relationships^2^.

Protein pre-clustering before signal triggering, or post-clustering upon receptor ligand binding has been extensively investigated^3–7^ and several microscopy methods have been developed to spatiotemporally resolve these events^8–12^. Super-resolution microscopy allowed disentangling nanoscale spatial and spatiotemporal events in living cells with unprecedented detail^13–20^. Collective lipid immiscibility in artificial and native membranes under equilibrium has been observed and its physical chemical and diffusional properties have been characterized extensively^21–27^. These studies have shown that collective interactions between sterols and saturated lipids induce the formation of condensed ‘liquid ordered’ (Lo) phases coexisting with ‘liquid disordered’ (Ld) phases, where the Lo phase displayed slower lateral diffusion. Heterogeneously distributed lipid nanodomains at the plasma membrane of cells, often referred to as rafts, are characterized by a dense lipid lateral packing, nanoscopic enrichment in sterols and saturated acyl chains, a transient lifetime, and a slow internal molecular diffusion^28–30^. Several methods have been developed to measure lipid diffusion^11,31–36^ to indirectly report on lipid nanodomain formation and protein diffusional hindering^37,38^.

Lipid heterogeneity is considered to play a crucial role in establishing communication platforms at the plasma membrane of cells, from where key proteins interact to prime their functions and trigger signaling events^38^. However, multidimensional spatiotemporal correlative imaging approaches of lipids at the plasma membrane are underexplored in living cells. Recently, by developing a spatiotemporal imaging approach that simultaneously records the same plasma membrane area in living cells at diffraction-limited and super-resolution, we reported that adaptive and lipid liquid-liquid immiscibility specialization and an active membrane remodeling is critical during primary cilium biogenesis^39^. An example of membrane specialization is the membrane of the primary cilium, which concentrates a large variety of receptors responsible for the transduction of extracellular cues by activating several signaling mechanisms^40–42^. The origin of the ciliary membrane depends on the route of primary cilium formation used^43–45^. In fibroblasts, the process of primary cilium formation starts intracellularly, and the ciliary membrane derives from a vesicle that progressively expands at the distal part of the mother centriole, and that then gradually deforms by elongation of an incipient axoneme whose distal end is encapsulated by a double membrane. Upon exocytosis in such a way that, upon its exocytosis, the most internal membrane becomes the ciliary membrane^45^. In contrast, in polarized epithelial cells, such as Madin-Darby canine kidney (MDCK) cells, which are a paradigm of renal tubular epithelial cells^42^, the process of assembly of the ciliary membrane takes place at the cell surface.

Our group reported that the midbody remnant (MBR), a structure resulting from the cleavage of the intercellular bridge formed during cytokinesis of MDCK cells moves along the apical membrane to meet the centrosome and enables primary cilium assembly^46^. More recently^39^, we have further investigated the role of the MBR and associated membranes in the assembly of the primary cilium specifically focusing on the role lipid immiscibility and collective dynamic self-assembly may play in ciliary membrane biogenesis. To this end, we developed a method that simultaneously resolves the spatiotemporal distribution of transient nanoscopic membrane subdomains and quantifies its molecular dynamics and lateral packing properties over time. Our study unveiled the role of the MBR, the source of lipids, and the role of lipid specialization and self-assembly in ciliary membrane biogenesis. Interestingly, by monitoring raft-like membrane dynamics in live cells using state-of-the-art super-resolution and fluorescence correlation spectroscopy-based techniques, we found out a membrane refinement process. A remnant-associated membrane patch (RAMP) spatiotemporally remodelled becoming enriched in more condensed membranes to form a centrosome-associated membrane patch (CAMP), which is the precursor of the ciliary membrane^39^. Employing our novel method based on laser interleaved confocal raster image correlation spectroscopy (RICS) and STED-RICS (LICSR)^39^, we unveiled multidimensionally and spatiotemporally lipid lateral packing and dynamics simultaneously. Strikingly, the refinement process before ciliogenesis showed micro- and nano- scale ordered lipid domains collectively organizing seemingly from hot spots of higher lipid condensation; however, we were unable to measure this collective organization in detail.

RICS and Number and Brightness (N&B) have been successfully applied independently to characterise the molecular dynamics of different cellular processes^47,48^ and its oligomeric state^2,49,50^. Indeed, we have shown that RICS and N&B can be combined to resolve spatiotemporally paxillin diffusion and oligomerisation state during the assembly and disassembly of focal adhesions^9^. RICS data can be correlated with N&B as the same set of images can be post-processed under certain conditions described elsewhere^9,51^. N&B yields a quantitative analysis of the relative number of molecules at different stages of oligomerisation^52,53^. Its combination with RICS can be limited by the spatial and temporal scale of the molecular signatures to resolve. For instance, N&B usually requires higher photon-counts, hence longer pixel dwell time, which renders its use to fast dynamics (e.g., cytosolic diffusion of low oligomeric state molecules) limited. Therefore, acquisition strategies shall be thought-through when correlating both techniques^9,51,54^. In this work, we have expanded our LICSR method and analysis and correlated it with N&B as depicted in the workflow showed in figure 1. By means of applying an N&B analysis to the same set of images recorded for LICSR, we quantified the relative number of molecules at the observed collective lipid hot spots spatiotemporally. This report aims at finding the sensitivity of this approach in unravelling lipid collective assembly, condensation, and dynamics multidimensionally and spatiotemporally, as well as to gain further insight into its possible limitations. Given its relevance in general cell biology processes and it particular in primary ciliogenesis, understanding how epithelial cells generate the ciliary membrane is a matter of fundamental biological importance and can broaden our understanding of lipid collective assembly in living cells.

**Figure 1.**
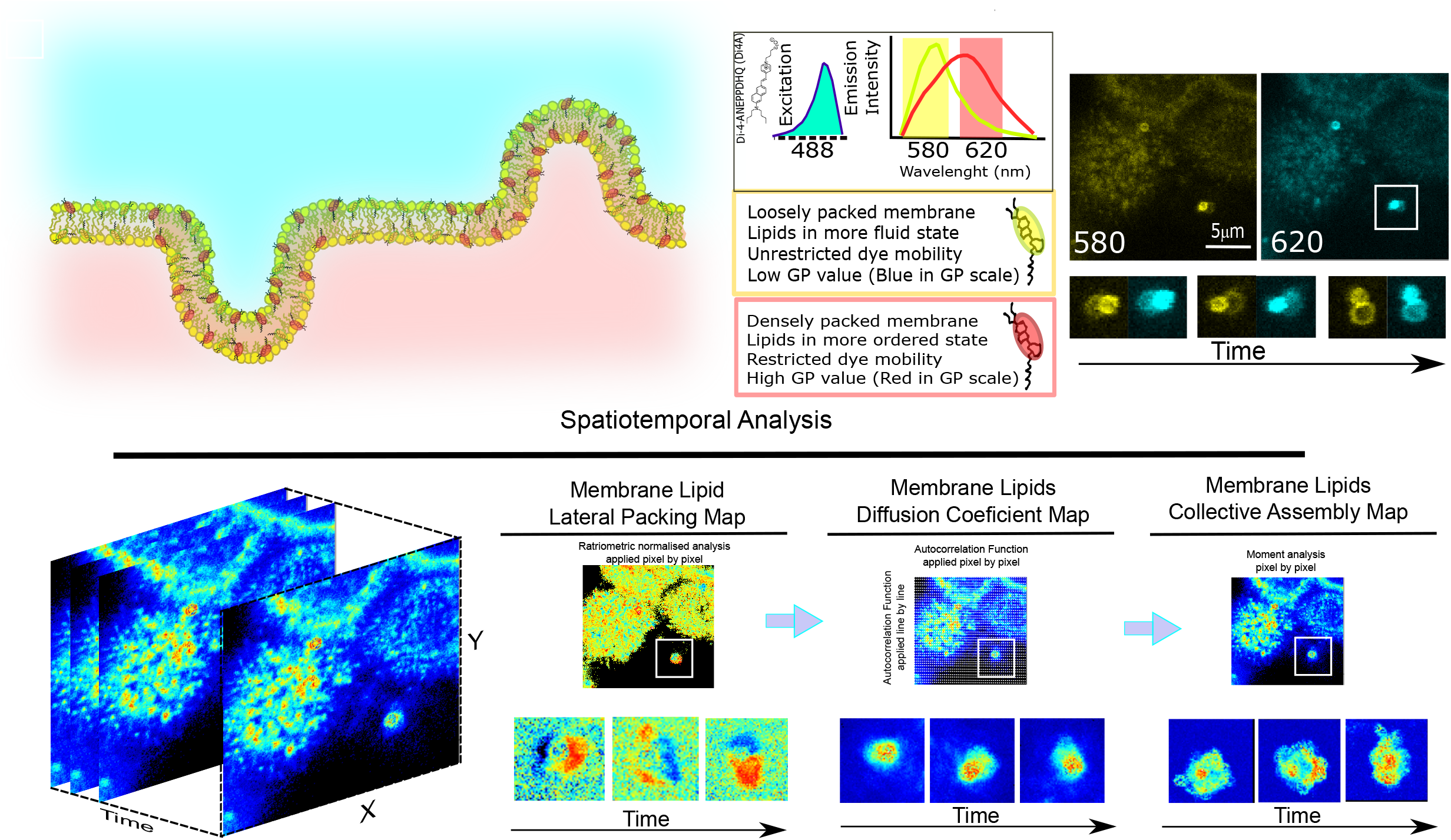
Experimental workflow for correlative LICSR N&B imaging with Di4A. Top row panel: from left to right: i) Representation of the membranes involved in this study, i.e., RAMP and initial stages of primary cilium genesis in epithelial polarised cells. ii) Scheme outlining the fluorescence properties of Di4A and the information that can be retrieved from its solvatochromic properties iii) Representative image of a MDCK cell labelled with Di4A showing the raw channels (560 and 620 nm, yellow and cyan respectively) to be ratiometrically normalised to quantify the GP index. Underneath the zoomed-in region highlighted inside a white solid frame square the images of both channels and how the RAMP/CAMP change over time. Bottom row panel: Scheme of the spatiotemporal correlative analysis employed in this work. Briefly, the recording of the same frame over time will be used to obtain the membrane lateral packing from the GP map; the same set of images will be used to obtain the membrane dynamics employing RICS, and it will be used to obtain the different diffusion modes by means of LICSR; employing the same set of images but applying the N&B analysis will inform about the membrane collective assembly properties. Overall, our multi-dimensional and spatiotemporal correlative imaging approach in live cells will be able to determine simultaneously lipid packing, its mobility at different spatial scales and its collective assembly over time at many spatial regions of interest.

## 2. Materials and methods

### 2.1 Cell lines and reagents

Epithelial canine MDCK II cells were obtained from the ATCC (ATCC® CRL2936) and cultured following provider’s instructions at 37°C and 5% CO_2_ in Modified Eagle’s Medium supplemented with 10% foetal bovine serum, 2 mM L-glutamine and antibiotics (1,000 U/ml penicillin and 0.1 mg/ml streptomycin) (all from Sigma-Aldrich, UK). Cellular membranes where labelled with di-4-ANEPPDHQ (Di4A) (ThermoFisher, UK).

### 2.2 Sample preparation, microscopy and STED imaging processing

MDCK cells were seeded on IbiTreat-coated 35mm dishes or μwell-chambers (both from Ibidi, #1.5) and grown for 4 days, until fully confluent. Cells were transferred to phenol-free media supplemented with 25 mM Hepes (Sigma-Aldrich/MERCK) and labelled with 40 μM Di4A prior to imaging. Di4A was added in excess to the culture media to allow constant labelling of the plasma membrane despite cellular internalisation. Imaging was performed at 37°C and 5% CO_2_ conditions.

Images were acquired using a Leica TCS SP8 3X gSTED SMD inverted confocal microscope (Leica Microsystems, Manheim, Germany) fitted with a HCX PL APO 63x/1.2NA CORR CS2 water immersion objective or a HC PL APO 100x/1.40 Oil STED WHITE. The excitation laser beam consisted of a pulsed (80MHz) super-continuum white light laser (WLL). Conditions have been previously described^9,39^; briefly, soluble cytosolic GFP for point spread function (PSF) calculation and Di4A were excited with at 488 nm; emission was detected between 500 and 550 nm for GFP and for Di4A two channels were recorded, 500-580 nm and 620 to 750nm. The STED depletion laser beam was a pulsed 775 nm; the emission was collected from 620 to 710 nm with a gating between 2 and 6.5 ns as in previous works^55,56^, and the pinhole was set to one Airy unit at 580nm. For a cleaner emission the excitation lines had a clean-up notch filter (NF) in the optical pathway. Images of 256×256 pixels at 16-bit depth were collected using 80.4 nm pixel size for confocal and 40.2 nm for STED and 8 μs pixel dwell time, for at least 200 consecutive frames. We measured the laser power at the back aperture of the objective. The average power of the WLL was 10-20 μW and the STED was 80-135 mW. gSTED nanoscopy, as expected, yielded images with low intensity counts. To maximise signal-to-noise we deconvolved the images using a commercially available software (Huygens package software, SVI, Netherlands). To quantify the background level of noise we used either an automated quantification provided by the software or a manual by means of computing the averaged background intensity from regions outside the object of interest or the cell. We obtained better results with the manual process. For the deconvolution we used 40 iterations, a signal to noise ratio of 15, and the classical maximum likelihood estimation method provided by the software. We characterised the resolution enhancement to be in average above 2-fold^39^. As in our previous paper^39^, the RICS and N&B analysis presented minimal differences when we used either the 500-580 nm or the 620-750 nm emission window. Since the most efficient depletion by STED was observed in the second, more red-shifted channel, the analysis of the data was performed in this channel. We found these conditions to be the best to minimize photobleaching while still being able to gather enough photons for the lipid order map imaging by means of the general polarisation index (GP), the membrane lateral diffusion calculations by RICS, and the collective lipid assembly by N&B.

### 2.3 Determination of the membrane lipid order by general polarisation parameter

The lipophilic polarity-sensitive membrane dye Di4A has been extensively used to characterise lipid membrane domains^57^. Like Laurdan^58^, it bears an electron-donor and electron-acceptor moiety to display large solvent-dependent fluorescence shifts. Di4A distributes at the membrane plane uniformly and ubiquitously, regardless of its lipid lateral packing properties (Figure 1). Its emission spectrum and location are independent of the phospholipid head groups and indirectly report the relative packing order or membrane condensation by sensing the local degree of water penetration. Most of these environmentally sensitive probes show an enhancement in charge separation when excited in polar solvents, which results in a larger dipolar moment. The penetration of water molecules into a bilayer composed of loosely packed lipids permits larger rotational and translational freedom to the probe, which yields an emission shift to longer wavelengths. Instead, when motion is more restricted due to sensing a milieu with lesser water content, the probe emission shifts toward shorter wavelengths. The ratiometric measure of the emission intensities at 580 and 620 nm yields the GP^59^. The GP allows the relative quantification of the membrane order or the degree of lipid lateral packing^60,61^. This method has been used to visualise native membrane microdomains in planar supported bilayers, giant unilamellar vesicles^62,63^ and in living cells^59,64–66^. The GP was calculated from fluorescence emission intensities collected at two wavelengths using the equation below:

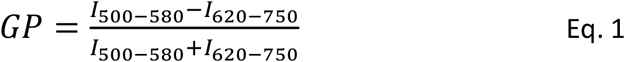

As previously described^39^, analyses were done for every pixel of the image using the commercial software SimFCS 4 (Globals Software, G-SOFT Inc., Champaign, Illinois). Similar analysis can be performed employing other software or open source macros^59,67,68^. Values of GP vary from 1 to −1, where higher numbers reflect lower fluidity or higher lateral lipid order, whereas lower numbers indicate an increased fluidity or lower lateral lipid order (Fig. 1). Analyses were performed over several cells (n>5). To improve the signal to noise ratio of the GP images, we averaged the GP at each pixel for 3 consecutive frames, and when the resulting image was highly pixelated, for presentation purposes only a 2×2 binning smoothing algorithm was employed.

### 2.4 Quantification of the molecular diffusion by raster image correlation spectroscopy

RICS uses the fluctuations of the fluorescence signal over time at a given PSF of molecules getting in and out an observation laser illuminated volume. RICS takes advantage of overlapping PSF analysis^9,69–73^. RICS analysis employs the image autocorrelation function on each frame (see Eq. 2):

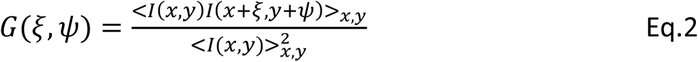

Where *I(x,y)* is the intensity at each pixel of the image, *ξ* and *ψ* are the spatial increments in the x and y directions, respectively, and *<I(x,y)>* is the average over all the spatial locations in both x and y directions of the image. The image correlation of all frames was averaged. The calculated spatial autocorrelation of the raster scanned pseudo image was then fitted in the x and y dimension with an equation that relates the correlation of the diffusion coefficient and the particle concentration in the illumination spot. The overall spatial autocorrelation function was given by:

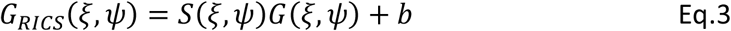

Where the spatial correlation *G*_*RICS*_(*ξ*, *ψ*) depends on the scanning optics *S*(*ξ*, *ψ*), the molecular diffusion correlation *G*(*ξ*, *ψ*) and the background (*b*), which for our study were as follows:

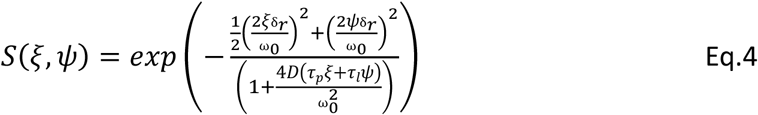

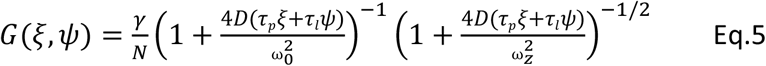

Where *N* is the number of particles in the PSF, γ is the illumination profile, which for a single photon excitation was considered 0.35^74^, *D* is the diffusion coefficient; *τ*_*p*_ the pixel dwell time; *τ*_*l*_ the time between lines; *δ*_*r*_ the pixel size; and *ω*_*0*_ and *ω*_*z*_ are the he beam waist in radial and axial direction. RICS advantageously can yield diffusion ranging from 0.1 to 1000 μm^2^/sec in an acquisition spanning from seconds to minutes; and permits a flexible selection of multiple regions of interest (ROIs) for post processing, and these ROIs can be analysed at different ranges of times within the whole stack of frames recorded, which enables determining spatial and temporal information tailored to the actual experimental investigated process. Determining more than one diffusion coefficient at the same ROI and at the same analysed timepoint is difficult, as they would need to be well-apart (e.g. typically more than one order of magnitude). However, employing different neighbouring ROIs and/or analysis the data during different sets of frames within the same ROI can reveal different diffusion coefficients^9,39^.

The PSF was characterised using purified recombinant EGFP (Biovision, Milpitas, California) diluted to a final concentration of 20 mM in PBS (Gibco, Thermo Fisher Scientific, Waltham, Massachusetts) or 1% BSA (Sigma-Aldrich/MERCK, Darmstadt, Germany) and transferred to a glass-bottomed μwell-chambers (Ibidi, Martinsried, Germany). Adjustments of the waist of the PSF were done by measuring the autocorrelation function of EGFP in solution with a fixed diffusion rate of 90 μm^2^/s^9,39,71^. We first did an estimation of the pixel dwell time to be used in the experiments by using the equation:

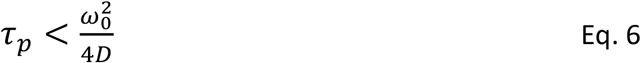

where, τ_p_ is the pixel dwell time; ω_0_, the waist of the PSF; and D, the diffusion of the analysed molecule. Two pixel dwell times were tested on cells traced with Di4A (4 and 8 μs) and we finally opted for 4 μs, as we found that at 256×256 pixel images and the pixels size of 80nm we could retrieve a good signal-to-noise per pixel that was compatible with the next method of analysis to correlate the images with, i.e., N&B. RICS analyses were performed using the commercial software SimFCS 4 (Globals Software, G-SOFT Inc., Champaign, Illinois). For RICS analysis we used a 2D autocorrelation function with no triplet state and a single diffusion component, also the 2D diffusion fitting routine was preferred instead of the 3D. A background subtraction filter of a moving average of 10 was used to discard possible artefacts due to cellular motion and slow-moving particles passing through, as previously detailed^9,39,75^. Analyses were performed over several cells (n>5).

### 2.5 Quantification of the spatiotemporal membrane diffusion modes by Laser Interleaved Confocal-STED RICS (LICSR) at multiple scales

LICSR^39^ was recently developed as an implementation of STED RICS^76^. LICSR employs a line interleaved laser excitation approach between Confocal and STED mode. By doing so, it enables, in a single measurement, resolving molecular diffusion modes at different spatial positions (two-dimensional i.e., X, Y) over time and at different spatial scales, i.e., nanometric and micrometric. As RICS, LICSR can map molecular diffusion being recorded in time-lapse mode from few seconds to several minutes. This is particularly useful to resolve in a single experiment different diffusion modes of lipids at the plasma membrane moving from 2D (x,t)^77^ to 3D (x,y,t) The spatial information allows direct visualisation of lipid micro and nano domains in real time (μs>t>s). Moreover, by means of labelling the plasma membrane with a photostable solvatochromic lipophilic dye, as for instance Di4A, LICSR deploys information about the lipid lateral packing at the molecular level in 3D (x,y,t). Taking advantage of the analyses we previously published for line interleaved scanning confocal and STED FCS^77^, we resolved in a single experiments in 3D (x,y,t) the different diffusion modes of lipids at the plasma membrane and the membrane packing properties.

To characterise the PSF for LICSR, as described above for RICS, 200 frames were recorded at 4 μs dwell time, i.e., 8.28 ms line repetition time (~1 sec per frame) of a 256×256 pixel region with a 20 to 40 nm pixel size of a EGFP solution (20 mM) and the waist of the PSF was calculated fixing the fitting to a known EGFP diffusion rate of 90 μm^2^/s^9,39,71^. The ratio between confocal excitation and STED power was optimised as previously described^39^ and we obtained a PSF of 234±7 nm for the confocal mode and 75±4 for the STED mode. For the live cell experiments, the optimal illumination efficiency and photon counts were achieved recording up to 200 frames with 40 nm pixel size at 8 μs. For the analysis and for comparative purposes, we obtained the diffusion coefficient of several regions of interest with the same size (32×32 pixels) at the same time-lapse. These considerations and settings were applied throughout the LICSR data acquisition and analysis. The analysis of the diffusion in confocal and STED mode was computed using the previously described fitting routines in the using the commercial software SimFCS 4 (Globals Software, G-SOFT Inc., Champaign, Illinois). Analyses were performed over several cells (n>5) To obtain the diffusion mode (i.e., Brownian, hindered,…) at which lipids move along the plasma membrane, we calculated the ratio between the diffusion coefficient in STED (D_STED_) and confocal (D_RICS_) mode^39,77^ (D_STED−RICS_), namely D_rat_. Quantification of D_rat_ at different ROIs and spatiotemporally permits to discern between free, trapped, and active diffusion. For instance, when D_rat_= 1, membrane diffusion is Brownian; however, when D_rat_<1 membrane diffusion is trapped, since its lipid molecular constituents’ diffusion is hindered because the dye is sensing the presence of highly condensed nanodomains. Trapped diffusion in condensed membranes resembles the previously described lipid diffusion in liquid ordered phases surrounded by lipids moving in a more fluid or liquid disordered phases; overall, forming liquid–liquid immiscible membranes^30,38,78,79^.

### 2.6 Quantification of the spatiotemporal membrane collective lipid assembly by N&B at multiple scales

N&B is a fluorescence fluctuation spectroscopy method capable of quantifying the average number and oligomeric state of freely diffusing molecules at each pixel of an image. This method takes advantage of measuring the brightness of a molecule or set of molecules counting the average of photons detected at a given time in the PSF. By employing moment analysis of the intensities detected over time per pixel and considering that the molecules are moving (i.e., not static, something that can be inferred from the RICS analysis performed previously), it can be calculated:

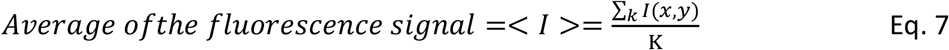

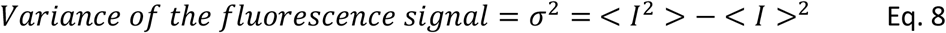

Where the sum is over the same pixel in each frame of the stack; *K* refers to the number of frames; *I* refers to the intensity measured in photon counts at one pixel of each frame; and the variance (*σ*^2^) is the amplitude of the fluorescence signal. From these, it can determined that at a given pixel

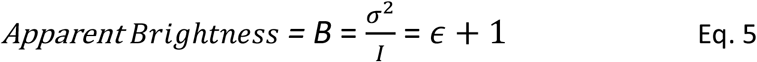

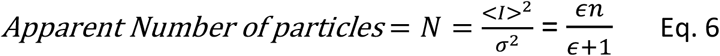

Where, as noted above, *B* the apparent brightness and *N* is the apparent number. Since the brightness of a fluorescent molecule (∊) is directly proportional to its oligomeric state (i.e., a tetramer would have four times the brightness of the monomer), it can be obtained the number of molecules per oligomer. Hence, the changes in brightness can be inferred directly with the fluorescence entity oligomeric state. The number of molecules (*n*) refer to an average-sized of the fluorescently labelled entity measured in counts per frame. This has been investigated for proteins so far and not yet for membrane lipids. Since lipids can collectively move and form transient domains with higher degree of lateral packing order due to chemical interactions and its thermodynamics^38,39^, we have employed this methodological approach to obtain direct visual and quantitative information of the collective assembly of lipids at the plasma membrane spatiotemporally. It shall be noted, however, that for immobile molecules B=1 and that the brightness provides a weighted average of those entities both immobile (should they occur over the stack of frames, B=1) and mobile (B=1+∊) with unknown weights^80^. Whilst finding and imaging functional immobile protein oligomerisation in live cell imaging is very complex, valuable information about changes in its oligomeric state can still be obtained: from low to high, or from high to low^9,80^. Since we are using this approach to investigate membrane lipids, the issue mentioned above does not interfere in the results because the lipids in living cells that may have lowest lateral diffusion are known to be in a (Lo)-like phases (often referred to as rafts).

N&B analysis was performed as described^50,53^ (Fig. 1). Postprocessing and quantifying of oligomeric state of proteins by N&B from pre-computed RICS data was done as previously reported^9^. We used the same set of time-lapse images from the LICSR data. To compute the brightness map, no detrending algorithm was used, as it has been described to be detrimental^80^. We obtained an average apparent brightness map and an apparent brightness versus an intensity scatter plot from the brightness analysis. The scatter plot served to yield a mirrored image of the average brightness map^9^. From this map, we spatially identified the collective lipid assembly in 2D (x,y). The apparent brightness values obtained were represented in occurrence histograms, which were then used to quantify the true number and brightness. Taking advantage of the quantification of the apparent brightness of free membrane lipid diffusion areas of the plasma membrane where low oligomeric state was observed, we could later resolve the degree of collective membrane lipid state. The apparent brightness was transformed to true brightness and the molecular brightness (cpms) subsequently calculated. These values served to identify multiples from the freely diffusing lipids (considered as a monomeric entity). These multiples were categorised in three groups: monomers/background, low collective membrane assembly, and high collective membrane assembly. The analyses of ROIs were performed similarly to those for RICS.

### 2.7 Quantification Software and Analysis

Statistical analyses and graph representation were performed using Origin2019 (OriginLab, USA). Results are presented as the mean ± standard deviation. Sample size and number of repeated experiments are stated in the legends. Statistically significant differences between pairs of data sets were analysed using the Bonferroni Test. All experiments had a minimum 3 biological replicates.

## 3 Results and Discussion

### 3.1 Condensed membranes associated to the MBR remodel at the nano- and micro-metric scale before ciliogenesis

The primary cilium is a highly condensed specialized plasma membrane protrusion harbouring many receptors involved in important signalling pathways^81^. Recently, we discovered that the source of the ciliary membrane in these cells is a remnant-associated membrane patch (RAMP)^39^. Once the RAMP reaches the centrosome area, it splits into two patches giving rise to the centrosome-associated membrane patch (CAMP), which ultimately feeds centrosome to assemble the ciliary membrane^39^. Super-resolution microscopy by means of STED nanoscopy (Fig. 2), 3D reconstruction approaches, and lipid diffusion quantification showed that membranes at the MBR/RAMP remodelled and stretched to form the nascent CAMP. This result is consistent with a previous a report showing that proper membrane condensation at the centrosome zone is necessary for efficient primary cilium assembly^82^.

**Figure 2.**
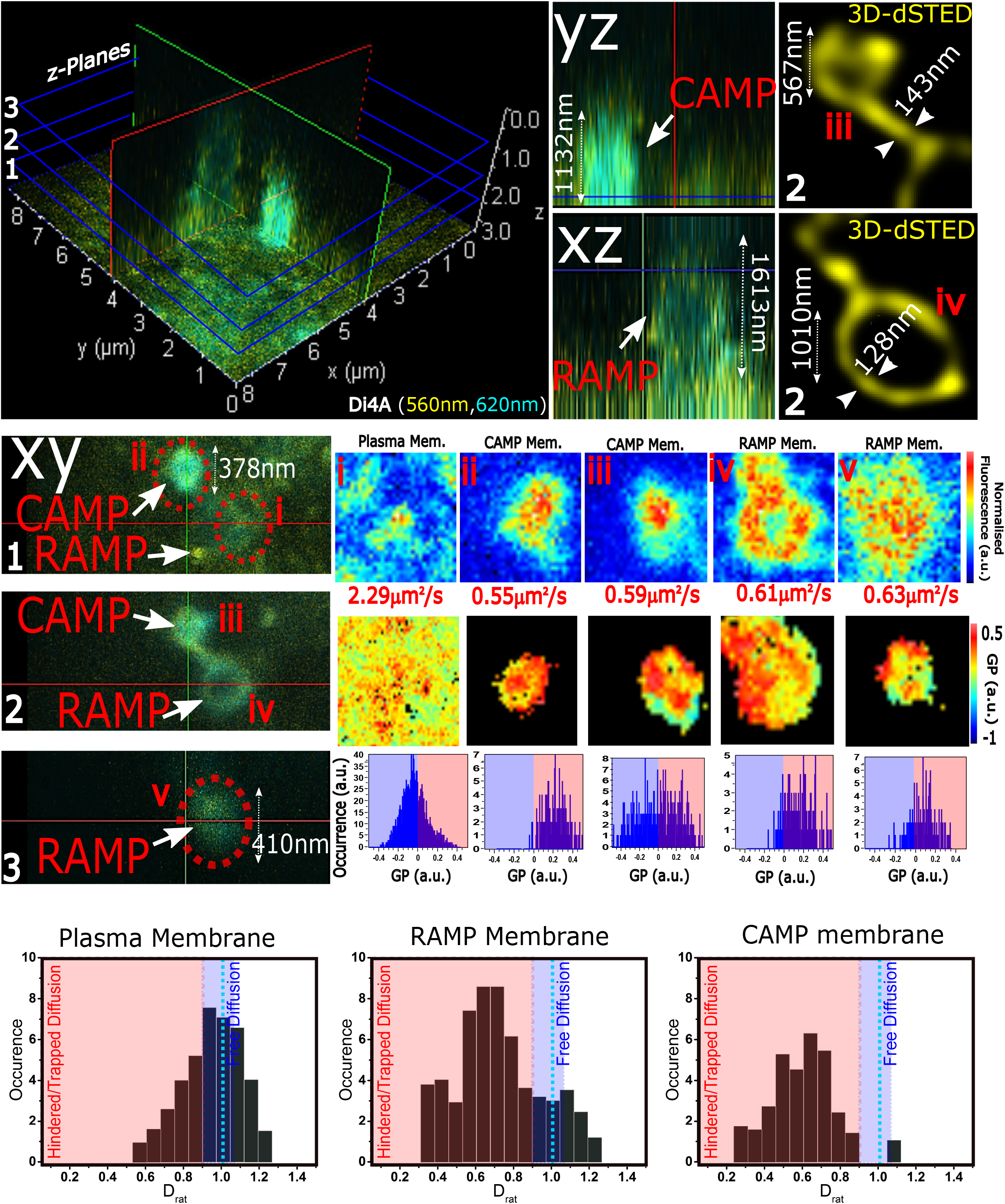
The RAMP has domains with condensed membrane and slow-moving constituents. MDCK cells showing a RAMP/CAMP labelled with Di4A. Top row panel, to the left-hand-side: large orthogonal axial reconstitution of raw micrograph view showing below at the central panel (left-hand-side) the XY section at 3 z-planes (z1: proximal, z2: middle and z3: distal); central images: YZ and XZ projections; right-hand-side images: zoomed-in deconvolved super-resolution STED image of the z2 middle plane. Central row panel to the right-hand-side from top to bottom row: diffusion coefficient values, intensity map and GP map of each of the cross sections z1-z3 and a plasma membrane region and occurrence histograms of GP values for cross sections z1– z3 and a plasma membrane region. Bottom row panel: frequency histograms obtained from LICSR showing the D_rat_ at the plasma membrane, RAMP and CAMP (>15 regions of interest from >7 cells). Red-shadowed region indicates hindered or trapped diffusion; blue-shadowed region indicates free diffusion.

In this report we have focused on the spatiotemporal remodelling process from the RAMP to the CAMP formation. To this end, we first determined membrane condensation by using the solvatochromic dye DiA4. We quantified the membrane diffusion properties by measuring the diffusion coefficient of the probe at the plasma membrane via RICS. The resulting membrane GP packing and molecular mobility maps confirmed the high level of condensation and slow diffusion of these specialised membranes (RAMP and CAMP) compared to its surrounding plasma membranes at the apical surface (Fig 2). The plasma membrane yielded a broad distribution of GP values. When fitting the histograms to a double gaussian fit we found a more fluid population of lipids, as observed through the low GP values (GP<0) and a population of lipids in a more ordered packing state, as observed with higher GP values (GP>0). However, resolving lipid packing at the nanoscale in live cells with this method was not possible. The statistical analysis of these populations showed an average coverage of condensed membranes of 41.29 ± 11.68%, with low GP values of −0.17 ± 0.12 and high GP values of 0.11 ± 0.17. Instead, the MBR/RAMP with overall similar GP values showed a microscopic lipid fluid and ordered distribution of 0.22 ± 0.17 (if present at all) and 0.34 ± 0.23, respectively, with a condensed membrane coverage of 58.18 ± 23.27% (Fig. 2). In conclusion, the results in Figure 2 illustrate a similarity in the degree of membrane condensation in RAMP and CAMP showing a microscopic heterogeneous distribution of ordered/disordered lipid packing areas, yet very different from that of the seemingly homogeneously lipid packing at the surrounding plasma membrane. RICS analysis of the Di4A at the membrane, yielded a diffusion coefficient that can directly inform about the lipid lateral mobility^39^. Since condensed but still fluid membranes retain the ability to allow lipid lateral diffusion^79^, we measured how membrane compaction correlated with molecular lateral mobility, although in a more restricted manner than in fluid membranes. We observed that the condensed membranes at the RAMP/CAMP had lower lipid lateral mobility at the microscale compared to the surrounded plasma membrane, 0.53 ± 0.2 and 2.31 ± 0.25, respectively (Fig 2), as expected.

The length and time scale resolution of RICS can be diminished by reducing the focal observation volume applying STED^76^. Thus, by combining STED with RICS (STED-RICS microscopy), we can assess the spatial molecular localization and the lipid lateral mobility at the nanoscale resolution (32). This method termed LICSR (laser interleaved confocal RICS and STED-RICS), yields the membrane lipid lateral packing and its dynamics at the micro and nanoscale simultaneously^39^. Applying LICSR to cells stained with Di4A resolves the diffusion and the physical-chemical state of the membrane in diffraction-limited (D_RICS_) and super-resolution (D_STED_) modes. Quantification of the ratio between them (D_STED_/D_RICS_, termed D_rat_) allows discrimination between free, hindered or trapped, and active diffusion^39,77^. Figure 2 shows the D_rat_ obtained from the plasma membrane, the RAMP and the CAMP membranes. Whilst the plasma membrane showed mostly membrane dynamics resembling a Brownian diffusion, RAMP and CAMP membranes yielded mobilities characteristic of those with a trapped molecular motion, most probably due to the high packing order of the lipids (i.e., raft-like nanodomains) at their membranes. Interestingly, the analysis showed a small fraction of lipids moving freely at the RAMP and, to a much lesser extent, at the CAMP. This is consistent with the results obtained with the GP maps, where regions of low and high lipid order within the RAMP and CAMP can be observed. The finding that RAMPs are highly condensed specialized membranes is consistent with previous studies showing that cells specifically regulate the localization of lipids to the midbody^83^, and with lipidomic analysis indicating that the midbody has a different lipid composition from that of most cellular membranes^84^.

### 3.2 The membrane at the RAMP remodels spatiotemporally adapting its lateral packing and molecular diffusion at the nano and micrometric scale

Once the RAMP is positioned in an apical central position, the newly formed CAMP remains at the plasma membrane zone above the centrosome. After separation of the patches, the MBR alongside the RAMP eventually disappear^39^. To better understand the process at the nanoscale the process of CAMP formation, we first acquired STED images of the RAMP and observed the presence of lateral segregation between ordered and disordered membrane regions within the same structure (Fig. 3). In addition, we tried to understand this phenomenon spatiotemporally. Figure 3 shows the continuous remodelling process that RAMP membranes undergo, where a clear reorganization of the membrane lateral segregation can be observed over time (i.e., interesting features upon formation and dissociation of budding membranes and ring-shaped structures). Overall, the structures become thinner as seen by a decrease of 48% in its perimeter size in average.

**Figure 3.**
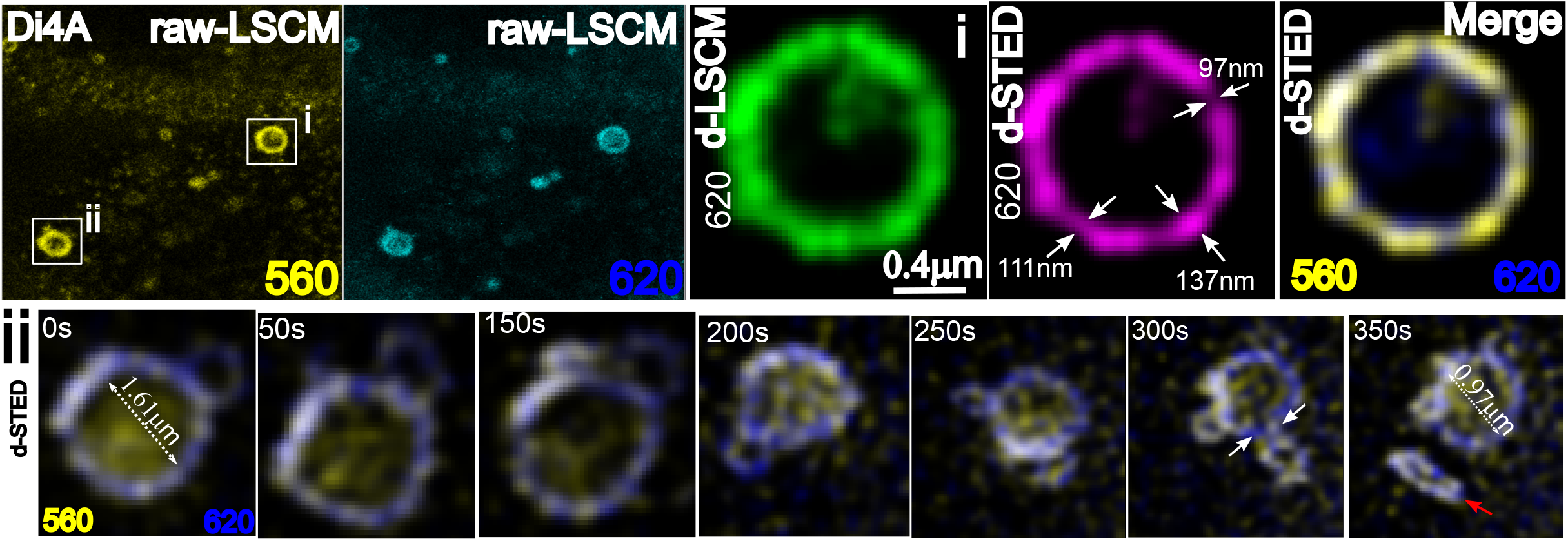
The RAMP remodels spatiotemporally by a refining process from heterogeneously distributed membranes. MDCK cells showing a RAMP/CAMP labelled with Di4A. Top row panel: raw laser scanning confocal microscopy images of the two Di4A emission channels (560 and 620 nm). Super resolution STED and laser scanning confocal microscopy (LSCM) deconvoluted images (both at from the 620 nm channel, green and magenta, respectively); merge of the deconvoluted LSCM (560 and 620 nm channels in yellow and blue respectively) showing the heterogeneous preferential spatial distribution of more condensed (560 nm channel) and less condensed (620 nm channel) Bottom row panel: Representative time-series of deconvolved LSCM micrographs (merged channels), where the temporal and spatial refinement process of the RAMP to form the CAMP can be observed. Of note, the refinement occurs by releasing smaller circular or elliptical-shaped membranes. White arrows and dashed lines indicate regions where size quantification was carried.

Finding of these packing coexistences led us to quantify the lipid lateral packing order by means of the GP index in space and time. The GP index could not be quantified at the nanoscale due to signal-to-noise issues and the ratiometric normalised values used. Common strategy to better observe changes are to either average several frames over time or to apply smoothing algorithms. We used an average of three frames to quantify the spatiotemporal distribution of the membrane condensation. The values obtained in Figure 4 increased over time while the RAMP perimeter decreased. This indicated a refinement in more condensed membranes over time. Noteworthy, the GP values ranged from 0.2 to 0.35 along the process; values that previously were described as (Lo)-like lipid phases^23,63^. The spatiotemporal evolution of the process revealed that the RAMP gradual reduction in size was due to continuous splitting of smaller ring-shaped membranes. These rearrangements result from an increased membrane tension at the connections or bridges between the larger and smaller structures. This process was mediated by an enhanced difference in the membrane packing properties at the necks of the budded structures (Fig. 4), which could be appreciated by the quantified difference between the high and low GP values (0.47<ΔGP<0.53), and consequently generated a membrane rupture and separation of the two patches^39^.

**Figure 4.**
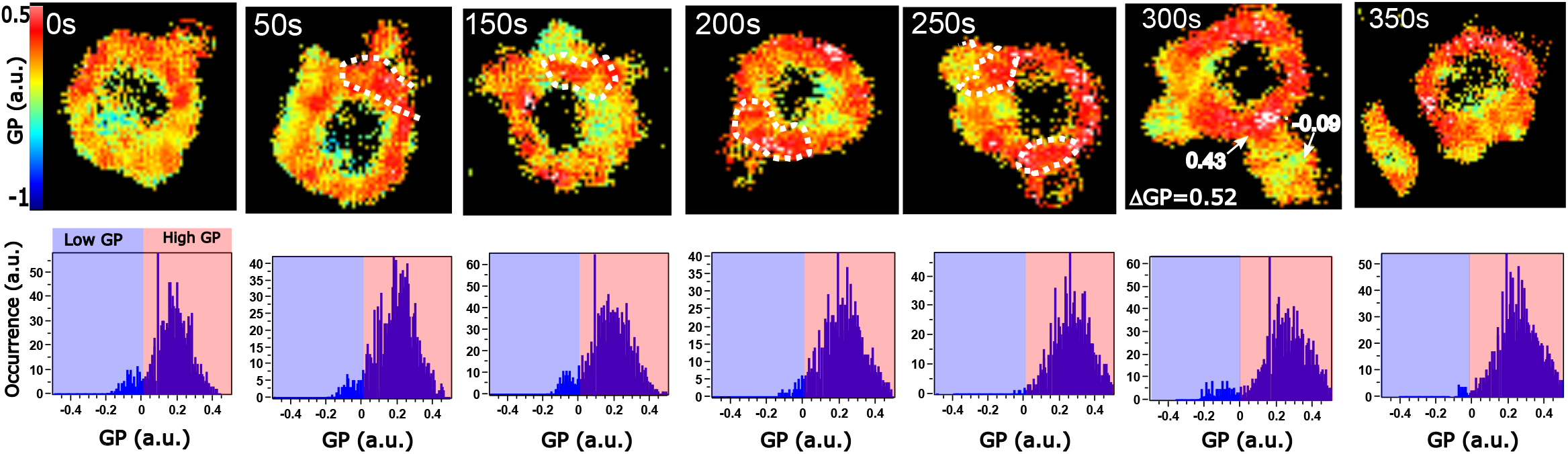
The RAMP remodels its heterogeneous distribution spatiotemporally and enriches in condensed membranes. MDCK cells showing a RAMP/CAMP labelled with Di4A. Top row panel: membrane condensation heterogeneous distribution in space and time is showed by the calculated GP maps, where pixels in cooler colours (blue to green) indicate more fluid membranes and pixels in warmer colours (yellow to red) indicate more ordered membranes. Dashed lines indicate the neck of the budding areas and number displayed the GP value and the difference in GP (ΔGP) between the budded area and the neck. The higher the ΔGP, the more differences in condensation, which is indicative of a more favourable scission by high difference in membrane tension. Bottom row panel: overall image GP frequency histograms. Blue-shadowed region indicates low GP values corresponding to fluid lateral packing or Ld-like membranes; Red-shadowed region indicates Lo-like membranes.

To better understand the evolution of the process with a higher spatial and temporal resolution, we applied LICSR. As mentioned above, the membrane dynamics and the lateral packing states correlate and give complimentary information about the physical-chemical properties of the membranes. The constituents of a condensed membrane move slower than in a more fluid membrane. Figure 5 shows how the diffusion coefficient decreased gradually with the structure refinement in condensed membranes. We observed a variation from 1 ± 0.4 to 0.09 ± 0.05 μm^2^/sec. A detailed quantification of the diffusion modes (D_rat_) of the probe at different regions of interest within the same membrane structure yielded always a characteristic trapped diffusion, as described elsewhere^34^. When the perimeter of the ring structure was reduced by >40%, the D_rat_ value was 0.4 ± 0.03, which indicated a highly hindered diffusion. In areas where a budding structure was present, the D_rat_ was slightly lower in line with previous findings, showing a large difference in the low and high GP values between structures (ΔGP). LICSR simultaneously resolved the spatial molecular localization, lipid lateral packing and the molecular diffusion at the micro- and nano-scale, determining the role of lipid immiscibility and heterogeneous distribution at the RAMP during its evolution towards the formation of the CAMP. These observations revealed highly valuable information about the structure-functional relationship in biology and the mechanisms involved in the physiological processes of membrane rearrangements^38^; however, the assessment of the nature, biophysical characteristics, and collective behaviour of membranes in real time is challenging.

**Figure 5.**
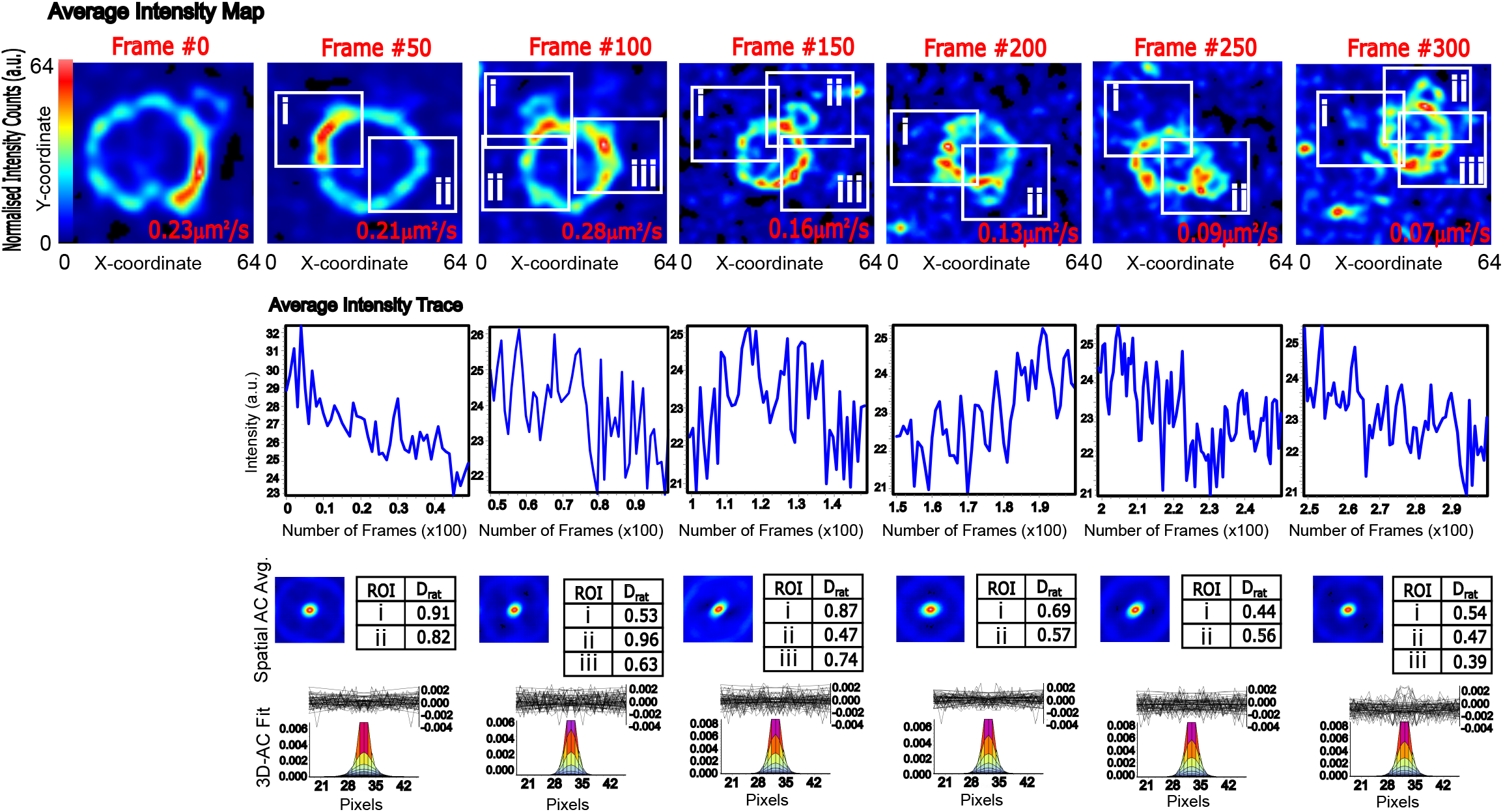
The constituents at the RAMP diffuse in a heterogenous fashion at the membrane spatiotemporally at different spatial scales. RICS analysis of MDCK cells showing a RAMP/CAMP labelled with Di4A. Analyses of a 64×64 pixel region of interest were done every 50 frames (300 total frames, 5-7 min). Top row panel: normalised average fluorescence intensity map of the RAMP/CAMP in space and time showing hot intensity spots that vary in its localization over time. At the bottom left corner of each image is shown the overall lateral Di4A diffusion coefficient at the membrane of the RAMP/CAMP. Middle row panel: average intensity trace of the 64×64 pixels region of interest variation over time showing no photobleaching. Bottom row panel: 2D average correlation maps and D_rat_ corresponding the white solid framed depicted regions of interests (ROI) at the top row panel. Lowest row panel show the autocorrelation fit of the STED RICS from the LICSR data represented in 2D of the obtained RICS analysis.

### 3.3 The spatiotemporal remodelling and adaptive condensation properties of the RAMP correlate with a collective rearrangement of its constituents

Our results highlight the importance of assessing liquid-liquid immiscibility remodelling and adaptive lipid condensation spatiotemporally. The high molecular order refinement and slow lateral diffusion, combined with the fact that these membranes are boundary-limited by a highly enriched protein region establishing a lateral diffusion barrier^85^, led us to think that in order to prepare the right scaffold for ciliary membrane biogenesis, these membranes should undergo spatial collective rearrangements. Hence, to further investigate if this process requires a modulation by means of a collective molecular assembly at particular hot spots of the CAMP, we decided to employ a multidimensional correlative imaging analysis of the LICSR data with N&B.

To this end, we had to consider that whilst these techniques are highly complementary, the analysis can be compromised by computing the images using different fluorescence intensity detrending algorithms, by applying different rolling image average or smoothing methods. For this purpose, we used raw images without any detrending or smoothing. However, to optimize the photon budget we used the same image average. For clarity reasons, images in Figure 6 are shown after the application of a 2×2 smoothing of the intensity.

**Figure 6.**
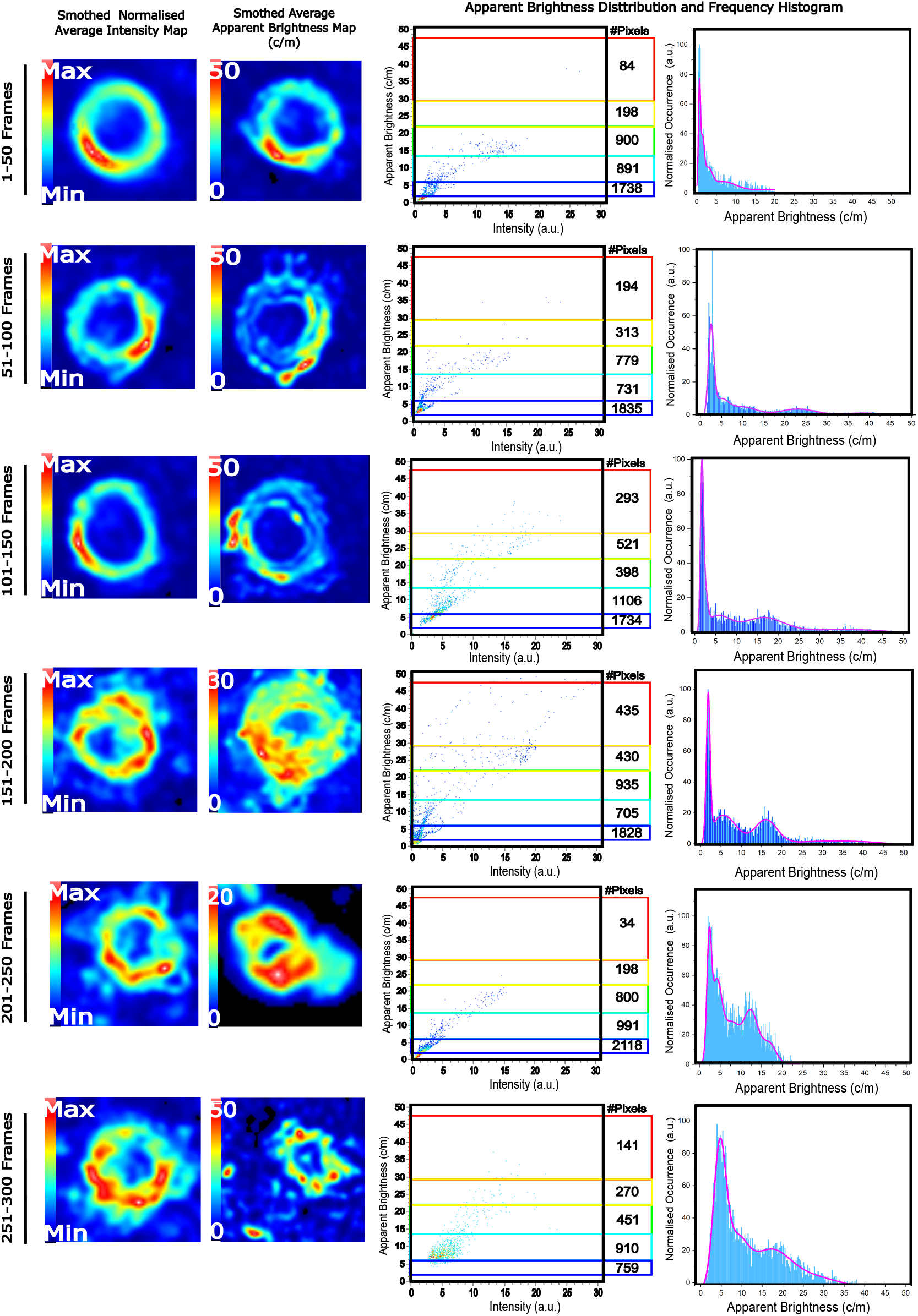
The constituents at the RAMP collectively concentrate and diffuse heterogeneously in space and time. N&B analysis of MDCK cells showing a RAMP/CAMP labelled with Di4A. Analysis of a 64×64 pixel region of interest were done every 50 frames (300 total frames, 5-7 min). Note that for correlative purposes, the same regions of interest and times as for the GP, RICS and LICSR were computed. Left-hand-side column panel: smoothed normalised average intensity map of the RAMP/CAMP at different times; next column to the right-hand-side shows the smoothed average apparent brightness map of the same region of interest. The following towards the right-hand-side column shows a pixel distribution map of the apparent brightness calculation from the N&B analysis. The color-coded rectangles correspond to the similar color code brightness map to facilitate the localization of the apparent brightness values in space and time. Inside the rectangles the value indicates the number of pixels with the corresponding apparent brightness. The uttermost column to the right-hand-side represent the frequency histogram of the obtained apparent brightness values at each image. N&B analysis reveal the spatiotemporal collective assembly of plasma membrane constituents at the RAMP/CAMP; the apparent brightness is a quantitative indication of the oligomerization or aggregation number of Di4A molecules.

An important aspect to consider is that N&B is more sensitive that LICSR to the movement of large objects. RICS-based methods minimise these possible artefacts by applying a filter, as described in the methods section. However, N&B large structures displacement can influence the precise localisation of the molecular oligomerisation under study, as it can be notice in some of the smoothed average apparent brightness in Figure 6. However, since we are interested in collective assemblies of lipids at the membrane, these methods can yield valuable information, as can be appreciated in the distribution and frequency histograms of the apparent brightness (Fig 6) The normalised occurrence clearly shows a population displacement towards higher apparent molecular brightness over time. At the first steps of the RAMP refinement, we could quantify most lipids in lower states (<10 c/m) of collective assembly. Interestingly but not surprisingly, the more refined the RAMP was, the more lipids shower higher collective assembly value (>10 c/m). Of note, when N&B was applied to protein oligomerisation determination, it could resolve tetramers, octamers and dodecamers^2,9,10,52,53^. Considering that on average in our STED PSF we could observed 93 ± 26 molecules of Di4A, as obtained from the single-molecule molecular brightness, one could estimate that the collective assemblies at the hot spots were formed by several groups of >10 c/m.

Taken together these results with the previous determination of the membrane molecular packing and the diffusion of molecules within the membrane, we can hypothesise that the refinement of the RAMP set the right scaffold to deliver highly specialised membranes from highly condensed lipid hot spots. This can open the way to better understand the importance of membrane sensing, self-assembly and remodelling in living cells, filling the experimental gap between membrane order and collective self-assembly.

## 4 Conclusions

During primary cilium membrane biogenesis occurs a molecular crowding favoured by an enrichment of the ciliary base in specialised lipids and proteins^39^. Certainly, the spatial and temporal molecular arrangements of these constituents play a critical role. However, revealing the functional role of the lipids has been elusive, mainly due to a lack of proper techniques able to quantify these events as they happen. The multi-dimensional and spatiotemporal correlative imaging carried out in this work in live cells at the plasma membrane has enabled us to determine the continuum nano-to-micro scale lipid adaptation and collective motion during ciliogenesis. Spatiotemporal molecular functional aggregation/oligomerisation is thought be involved in many physiological mechanisms, where lipid liquid-liquid immiscibility has been placed at the spotlight^38^. Notably, we have revealed that the RAMP refinement is facilitated by a continuous membrane condensation with lipids becoming gradually more entrapped and moving orchestrated in a collective manner. Membrane condensation is highly dependent on collective protein enrichment/oligomerisation and lipid self-assembly. Lipid phases have been shown to rely on molecular lateral packing, membrane lateral pressure, and crowding. It is believed that these molecular enrichment and specialisation enables biochemical reactions and infers physical viscoelastic properties^38^. Membrane lateral diffusion and order, recently, could be determined simultaneously; however, no collective molecular insights to indirectly provide self-assembly spatiotemporal information could be provided. Here, combining LICSR and N&B, we have added that extra important layer of quantitative data. This work led us to hypothesise that together with the known important role of certain individual lipids (i.e., cholesterol, phosphatidyl serine, inositols, etc) and the existence of lipid immiscibility in living cells, we should also consider membrane phase-independent, probably protein-aided, collective assembly as a functional determinant in cell biology.

Renal cystic diseases are the most common of the many abnormalities associated with ciliary dysfunction^86^, making research on the primary cilium of renal epithelial cells particularly important. Our study^39^ identified the source of the ciliary membrane of cells, such as that of renal epithelia, whose primary cilia are assembled entirely at the plasma membrane by revealing that the ciliary membrane arises from a patch of condensed membranes that the MBR delivers to the centrosome. The new method that we describe in this work, that combines LICSR and Number and Brightness (N&B) allowed us quantitative determine at different spatial scales (micrometer to nanometer) and over time (from seconds to minutes) the membrane lateral packing properties, its diffusion mode and molecular mobility and whether the process of CAMP formation from the RAMP involves a lipid collective assembly in form of membrane hot spots; overall, providing novel insights about the important role collective lipid assembly at the plasma membrane may play spatiotemporally in cell biology. These findings revealed the mechanism of biogenesis of the CAMP, which is the precursor of ciliary membrane of polarized epithelial cells and provide novel insights into how post-mitotic midbodies influence cell function.

## ACKNOWLEDGEMENTS

This work was supported by funding from a Marie Curie Career Integration Grant (NanodynacTCELLvation; PCIG13-GA-2013-618914) to JBS and a grant (PGC2018-095643-B-I00) to MAA from the Spanish Ministerio de Ciencia e Innovación, Agencia Estatal de Investigación, y Fondo Europeo de Desarrollo Regional, European Union (MICINN/AEI/FEDER, EU) and by the financial support of the Central Laser Facility (Science and Technology Facilities Council, Harwell, UK) that enabled us to use in-house confocal and STED equipment under application number 16230026. The authors declare no competing interests

## Notes

### Competing Interest Statement

The authors have declared no competing interest.

